# Biological invasion drives ecosystem state and metabolism across tipping points

**DOI:** 10.64898/2026.04.16.718977

**Authors:** Emilien Gaigné, Olivier Dézerald, Caroline Gorzerino, Julie Coudreuse, Yoann Bennevault, Alexandrine Pannard, Eric Edeline

## Abstract

Abrupt regime shifts of complex ecosystems between alternative stable states are widespread in nature. Yet, our mechanistic understanding of disturbance-shift-ecosystem functioning relationships remains poor, and it is further unclear whether biotic disturbances can drive such shifts. Using a 5-year pond experiment, we demonstrate that invasion by the red swamp crayfish (*Procambarus clarkii*) drove a regime shift from a clear-water, macrophyte-dominated, to a turbid, phytoplankton-dominated state. The regime shift was associated with increased water temperature due to increased water turbidity enhanced light absorption, and with a seasonal switch of ecosystem metabolism from hetero-to autotrophy due to decreased respiration in summer, despite constant gross primary production. Reducing crayfish population densities by 44 % failed to move ecosystems back towards their initial state and functioning. Our results stress that biotic disturbances may have hardly-reversible consequences on the biophysical and biogeochemical processes that support ecosystem functioning.

## Introduction

Ecosystem responses to disturbances, be they natural or anthropogenic, can lead to hardly-reversible shifts between alternative stable states when an environmental condition is changed beyond a threshold value, or a ‘tipping point’ (Beisner *et al*. 2003; Scheffer *et al*. 2001). Such drastic regime shifts, documented in terrestrial, marine and freshwater ecosystems (Done et al. 1992; Hare & Mantua 2000; Tucker & Nicholson 1999), often entail major ecological and socio-economic consequences, including species loss or declines in ecosystem services (e.g. reduced water quality; Janssen *et al*. 2021; Kéfi *et al*. 2016). Yet, despite several decades of research, our understanding of the mechanisms underlying the maintenance of multiple stable states and the transitions between them in complex ecosystems remains limited (Kéfi *et al*. 2022). This gap partly stems from the predominance of empirical research relying mainly on circumstantial observations (e.g. Capon *et al*. 2015; Higgins *et al*. 2024; Mata *et al*. 2025), whereas understanding the mechanistic links between disturbances and regime shifts requires long-term manipulative experiments (Scheffer & Carpenter 2003; Schröder *et al*. 2005).

Shallow lakes and ponds provide the best-supported empirical example of complex ecosystems subject to bistability. The two alternative stable states have been defined as (1) a clear water state dominated by submerged macrophytes, and (2) a turbid state dominated by phytoplankton (Scheffer *et al*. 1997). The bistability is commonly attributed to self-reinforcing biotic and abiotic feedback loops involving interactions between submerged vegetation, phytoplankton and turbidity (Scheffer *et al*. 1993, Kéfi *et al*. 2016), which may result in hysteresis, a form of abrupt bistability where reverse transitions occur at different disturbance levels (Schröder *et al*. 2005). In the clear-water state, macrophytes prevent the invasion of phytoplankton by monopolizing nutrients, and by hindering water mixing and nutrient resuspension from sediments. Abiotic disturbances, mainly through increased nutrient loading, can negate the competitive advantage of macrophytes over phytoplankton, and push shallow lakes to a turbid state in which macrophytes can no longer persist or invade (Scheffer *et al*. 1993, Kéfi *et al*. 2016).

In contrast with abiotic disturbances, the roles of non-human biotic disturbances as drivers of ecosystem regime shifts have received comparatively little attention (Kéfi *et al*. 2016; Mata *et al*. 2025; Su *et al*. 2021). In particular, although strong interactors such as megafauna or invasive species can profoundly affect the structure and function of freshwater, marine and terrestrial ecosystems (Ehrenfeld 2010; Estes *et al*. 2016; He *et al*. 2024; Malhi *et al*. 2016), a pervasive view remains that biotic disturbances are not strong enough to trigger regime shifts in complex ecosystems, and can only modulate the tipping points of ecosystem collapse and recovery (Bauer 2012; Diamond *et al*. 2022; Reynolds & Aldridge 2021; Rocha *et al*. 2015). This view, however, largely relies on either short-term experiments or on circumstantial empirical observations in which the effects of biotic drivers are confounded with the concomitant effects of multiple other environmental drivers. To fill this knowledge gap, our first objective was to provide first long-term experimental evidence that a biotic disturbance can trigger a regime shift in freshwater pond ecosystems. As biotic disturbance, we use the invasive red swamp crayfish *Procambarus clarkii*, a species assumed to be capable of driving a regime shift of clear-water ponds to a turbid state based on field observations (Matsuzaki *et al*. 2009; Rodríguez *et al*. 2003).

Another major gap in our understanding of regime shifts relates to their consequences on ecosystem functioning (Flood *et al*. 2020; Hilt *et al*. 2017), defined as the processes governing fluxes of energy and materials (Strayer 2012). Ecosystem metabolism is the balance between gross primary production (GPP) and ecosystem respiration (ER) yielding net ecosystem production (NEP = GPP – ER). NEP is thus an integrative indicator of whole-ecosystem functioning and integrity (Jankowski *et al*. 2021) that determines whether an ecosystem is predominantly heterotrophic (NEP < 0; i.e., a net carbon source) or autotrophic (NEP > 0; i.e., a net carbon sink). In freshwater ecosystems, shift from a macrophyte to a phytoplankton state driven by eutrophication has been associated with increased NEP due to decreased ER and stable GPP (Brothers *et al*. 2013, Diamond *et al*. 2022). However, at the same time, turbid phytoplankton-dominated lakes and ponds may be warmer due to increased light absorption (Paaijmans *et al*. 2008, Torma & Wu 2019), and warmer systems are expected to have decreased NEP, because ER increases at a faster rate than GPP with warming (O’Connor *et al*. 2009; Yvon-Durocher *et al*. 2010). These two conflicting effects of shifts in dominance of primary producers and temperature on NEP might explain frequent inconsistencies in the empirically-observed effects of regime shifts on lakes and ponds metabolism (Hilt et al. 2017). To help clarifying these mechanisms, our second objective was to use our long-term experiment to quantify the effects of a crayfish-induced ecosystem regime shift on both NEP and temperature at the same time.

To simultaneously address these knowledge gaps, we used a five-year manipulative experiment in freshwater outdoor mesocosms, emulating vegetated pond ecosystems. Crayfish were introduced more than 1.5 years after pond establishment and, 2 years after crayfish introduction, we added a crayfish-removal treatment that reduced crayfish density by 44% on average. Crayfish removal aimed at testing for nonlinearity in the disturbance-shift relationship, which is a hallmark of alternative stable states. During the whole experiment, we monitored macrophyte cover, phytoplankton concentration and water turbidity and, during the last year, we additionally measured water temperature and oxygen concentration at 10-min intervals, from which we reconstructed ecosystem NEP, GPP and ER.

We predicted bioturbation and macrophyte grazing by crayfish to drive a regime shift from a macrophyte-dominated, clear-water state to a phytoplankton-dominated, turbid state (Matsuzaki et al. 2009; Rodríguez et al. 2003, Reynolds & Aldridge 2021). We further predicted the regime shift to both decrease ER (Brothers *et al*. 2013) and to increase water temperature at the same time (Paaijmans *et al*. 2008, Torma & Wu 2019). Then, depending on the relative importance of changes in primary producers (Brothers *et al*. 2013) and in temperatures (O’Connor *et al*. 2009; Yvon-Durocher *et al*. 2010), we expected the NEP to either increase or decrease, respectively. Finally, because shallow lakes and ponds are subject to hysteresis (Scheffer *et al*. 1993, Kéfi *et al*. 2016), we expected disturbance relaxation from crayfish removal to be insufficient to restore the uninvaded ecosystem state, but to be possibly strong enough to move ecosystems into the bistability region where the two alternative states coexist.

## Materials and Methods

### 1. Study design

#### Model species

*P. clarkii* is native to the southern United States and is considered one of the most widely introduced freshwater species in the world, especially due to its high economic importance (Souty-Grosset *et al*. 2016). This crayfish is tolerant to changes in environmental conditions, including cold and drought, and often has major impacts on invaded ecosystems through its burrowing and foraging activities (Twardochleb *et al*. 2013).

#### Experimental design

In October 2019, twelve 11-m^2^ pond mesocosms (7,000 L) were installed at the INRAE experimental facility for aquatic ecology and ecotoxicology in Rennes (France, PEARL; https://www6.rennes.inrae.fr/u3e_eng/). Mesocosms received a 5-cm layer of pond sediments mixed at 50% with river sand and they were filled with tap water. Mesocosms were also covered with 10-cm meshed nets that prevented avian predation, but that allowed a colonization by other organisms (e.g. large flying macroinvertebrates). From both the seed bank and *de novo* colonization, macrophytes and a complex pond community rapidly developed. Additionally, we introduced the same number of *Lymnaea stagnalis* in each mesocosm since gastropods were not naturally present (N = 45 adults in March 2020; N = 421 juveniles in July 2024). By spring 2020, all ponds hosted a fully developed and stable macrophyte community, dominated by *Chara globularis* and *Alisma plantago aquatica* or filamentous algae, and were all in a macrophyte-dominated, clear-water state (Fig. S1A). In our analyses, filamentous algae were included in the ‘macrophytes’ category because, similar to vascular plants, they outcompete phytoplankton for light and nutrients by forming large mats.

In June 2021, 40 crayfish were introduced in each of eight mesocosms (20 males and 20 females). Crayfish originated at 50% from two independent populations: Brière Marsh (western France) and Lake Lamartine (South-West France). At crayfish introduction, we also added in each mesocosm twelve artificial refuge traps (ART) that provided crayfish with shelters (Edeline *et al*. 2025; Green *et al*. 2018). ARTs are made of PVC tubes glued together and closed by a grid at one end. Adding ARTs also in uninvaded mesocosms ensured that crayfish effects were decoupled from ART effects. We tracked crayfish population dynamics using mark-recapture. Crayfish were trapped using ARTs and commercial traps (metal, 1-cm meshed, see Fig. S2) and were marked using either PIT tags (total length > 54 mm) or coloured nail polish (20 mm < total length ≤ 54 mm). As a proxy for total crayfish population size (Fig. 1), we used the number of uniquely-captured individuals during a mark-recapture session, which lasted for 5 consecutive days, repeated twice per year.

**Figure 1:**
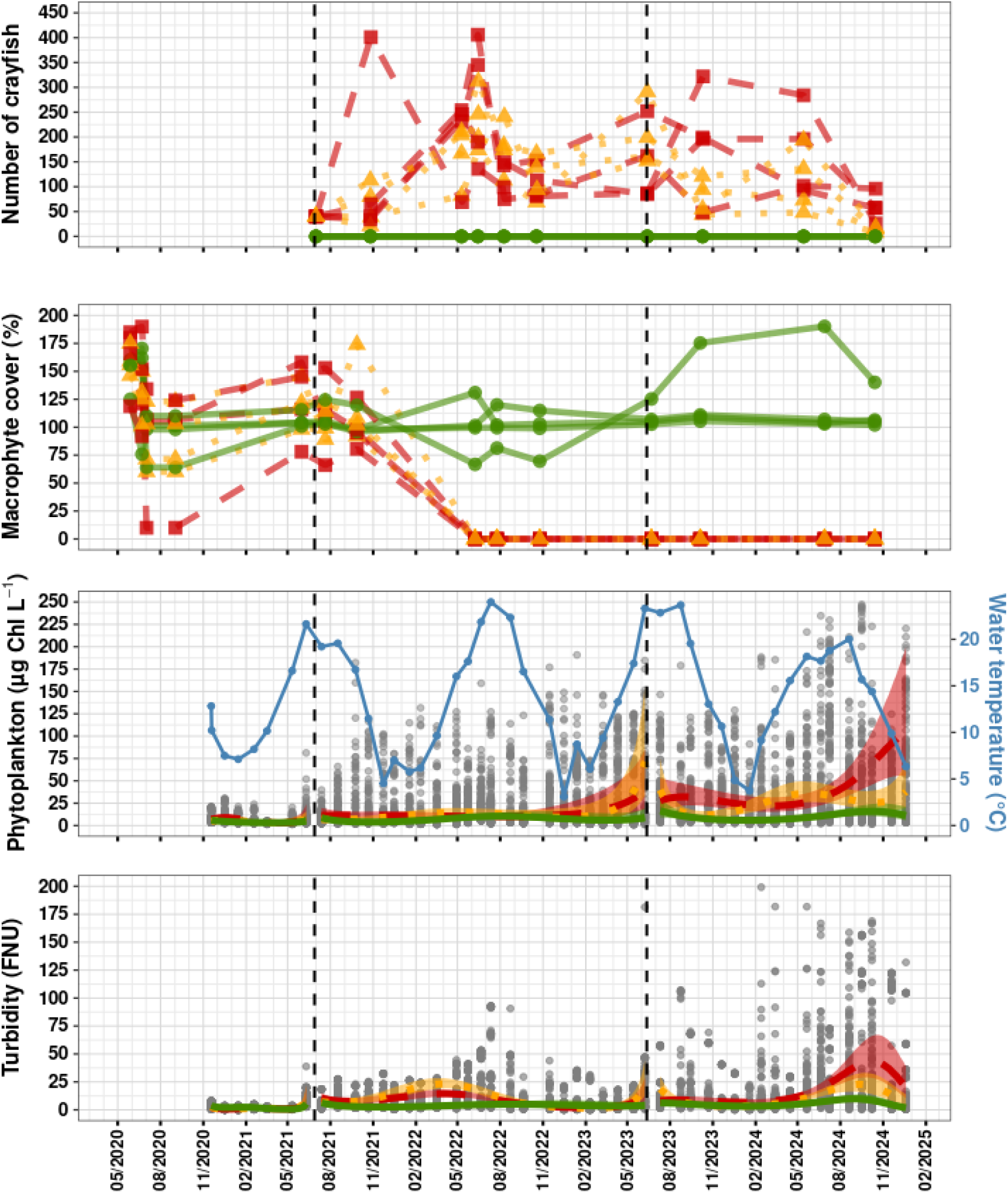
Invasion impacts on ecosystem state. Lines represent the number of crayfish, percent cover of macrophyte and mean water temperatures, or predictions from LMMs with 95% confidence intervals for phytoplankton concentration and turbidity. Line types reflect the experimental design: uninvaded control (green solid line), invaded (red dashed line), and harvested mesocosms (orange dotted line). For modelling purposes, treatments were assigned to mesocosms prior to their effective implementation. Vertical dashed lines indicate the timing of crayfish introduction and harvesting in July 2021 and in June 2023, respectively. Grey dots show the raw data. For improved visualization, some extreme values are not displayed, but were included in the models (phytoplankton: N = 45; turbidity: N = 33).

From June 2023, four of the eight crayfish populations have been harvested annually, resulting, on average, in a 44% decrease in crayfish densities (Fig. 1). The objective was to test for the ability of the ecosystems to recover their initial, uninvaded state. Thus, the experiment was split into three distinct periods: Period 1 of fully-developed, uninvaded communities (1^st^ January 2020 - 1^st^ July 2021, 12 uninvaded mesocosms); Period 2 after crayfish invasion but before harvesting (1^st^ July 2021 - 12^th^ June 2023, 4 uninvaded vs 8 invaded mesocosms), and Period 3 during crayfish harvesting (12^th^ June 2023 - 20^th^ December 2024, 4 uninvaded *vs* 4 invaded *vs* 4 harvested mesocosms).

#### Ecosystem-state measurements

Macrophyte communities were monitored three times per year for species composition and species-specific percentage cover (visually assessed) by a botanist. Turbidity (Formazin Nephelometric Units, FNU), total chlorophyll concentration (µg Chl L^-1^; i.e., a proxy for phytoplankton concentration), and water temperature (°C) were measured monthly using a multiparameter water quality probe (YSI ProDSS, Turbidity sensor 626901, YSI-626210 probe). Measurements were made at a depth of 10-20 cm, at twelve locations evenly distributed across the mesocosm surfaces. We excluded from the turbidity data one negative value and 11 implausibly high values (>1000 FNU).

### 2. Metabolism estimation

From 26^th^ September 2023 to 10^th^ October 2024 (Period 3; Fig. 1), dissolved oxygen (DO; mg L^-1^) and water temperature (T; °C) were recorded every ten minutes in each mesocosm using high-frequency sensors (HOBO-U26-001) positioned approximately 30 cm above the bottom. Probes were cleaned every two weeks to prevent fouling. Every three months, data were downloaded, and probes were installed in 100% air-saturated water to check for drift over time. These checks confirmed that sensor drift remained negligible throughout the study period and that all sensors provided consistent values.

We then estimated daily rates of gross primary production (GPP; mg DO L^-1^ d^-1^), ecosystem respiration (ER; mg DO L^-1^ d^-1^), and net ecosystem production (NEP = GPP – ER) using the diel oxygen technique (Staehr *et al*. 2010), implemented in the package ‘Lake Metabolizer’ (Winslow *et al*. 2016), which the detailed approach is provided in Supplementary Material (Appendix 1). Briefly, changes in dissolved oxygen concentration (DO) were modeled over 10-minute intervals (t) using the following equation:

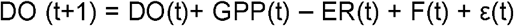

where F(t) represents air-water gas exchange, and ε(t) a normally-distributed process error.

Estimating ecosystem metabolism requires meteorological data. We thus collected shortwave radiation (W.m^-2^), atmospheric pressure (hPa), and wind speed (m.s^-1^) every five minutes from a nearby meteorological station located within 100 meters from the mesocosms (meteorological station of the Institut Agro; https://geosas.fr/stav/vidae-2-accueil.html). These data were linearly interpolated to match the 10-minute resolution of temperature-oxygen sensor data.

### 3. Statistical analysis

To quantify the effects of the three experimental treatments (uninvaded, harvested, invaded) on macrophyte cover, phytoplankton concentration, turbidity, water temperature, and ecosystem metabolism (GPP, ER, and NEP), we fitted linear mixed-effects models (LMMs), assuming Gaussian distributions. Skewed positive variables (turbidity, phytoplankton concentration) were natural log-transformed prior to analysis.

In all LMMs, fixed effects included experimental Treatment (factor), Time, and Treatment-by-Time interactions. Time was modeled either as a Period (factor) or as a polynomial term of Day or Day-within-Period effects, depending on the temporal resolution of the data. We used third- or fourth-order polynomials depending on goodness of fit, as visually-evaluated from plots of residuals vs. fitted values. Both Day or Day-within-Period were standardized to zero mean to reduce spurious slope-intercept correlations.

For macrophytes, we fitted single model with a period factor because the low number of measurements per period precluded meaningful characterization of within-period temporal dynamics. For phytoplankton concentration and turbidity, we fitted one separate model for each experimental period, each including a Day-within-Period; hence resulting in three models for phytoplankton concentration and three models for turbidity. For water temperature and ecosystem metabolism (GPP, ER, and NEP), we fitted one separate model to each variable using Day as the Time effect, as these measurements were conducted only during a part of the Period 3 (see Tables S1 and S2 for complete models).

Importantly, in modelling macrophyte cover, phytoplankton concentration and turbidity, we assigned mesocosms to their experimental treatment from Period 1 onwards. Although this coding does not reflect the reality of the experimental design (treatments were added sequentially across periods), it allowed us to test for possible unexpected prior-to-treatment differences attributable to random mesocosm effects during Periods 1 or 2. A drawback of this coding is that it generated spurious Treatment-by-Period interactions, which we therefore did not interpret.

In total, we fitted eleven LMMs, all of which included mesocosm identity as a normally-distributed random effect on both the intercept and temporal dynamics. Model assumptions were verified through residual diagnostics, including checks for normality, homoscedasticity, and independence using the ‘performance’ R package (Lüdecke *et al*. 2021). When present, residual heteroscedasticity was modelled by making the log of residual variance linearly dependent on experimental treatments and periods and their interaction in LMMs (“Dispersion” sections in Table S2). All analyses were conducted using Rv.4.x (R Core Team 2025) with models fitted using the ‘glmmTMB’ R package (Brooks *et al*. 2017).

## Results

### 1. Impact of invasion on ecosystem state

Deviance analyses for each of the seven LMMs for macrophytes, phytoplankton and turbidity dynamics are provided in Table S1. Overall, all dynamics were significantly influenced by the experimental Treatments, and by a Treatment-by-Time interaction, except during the Period 1 when the treatments were not experimentally implemented. Deviance analyses of LMMs for phytoplankton and turbidity further revealed that there were significant seasonal dynamics that were influenced by the treatments (significant « DAY_period^n^ », and « Treatment × DAY_period^n^» effects in models 2-7 in Table S1).

Hereafter, we used ‘Treatment’ as an umbrella term for both Invaded and Harvested effects in Table S2. We now focus on each experimental period separately. During experimental Period 1 corresponding to the 20 months that followed mesocosm installations, and prior to crayfish invasion, ecosystems were in a clear-water, macrophyte-dominated state (Fig. S1A), which was characterized by a high macrophyte cover, low phytoplankton biomass and low turbidity (Fig. 1; Fig. S1A). Macrophyte colonization was remarkably fast, since all ponds reached more than 100% cover by May 2020 (ca. seven months after the mesocosm installations). The drop in macrophyte cover in July 2020 was due to disappearance of filamentous algae that initially dominated some mesocosms (Fig. 1). After June 2020, filamentous algae were no longer recorded in any mesocosms. Importantly, during experimental Period 1 there were no significant pre-existing differences in macrophyte cover, phytoplankton biomass or turbidity among mesocosms across experimental treatments (Fig. 1, non-significant « Treatment » effects in models 1, 2 and 5 Table S2).

During Period 2 corresponding to the 23 months following *P. clarkii* introduction, macrophyte cover dropped in invaded mesocosms while remaining stable in control ones (Fig. 1, Fig. S1B, significant « Treatment × Period 2» effects in model 1 Table S2). Ten months after crayfish introduction (May 2022), macrophytes were completely lost from all invaded mesocosms (Fig. 1). In parallel, crayfish increased both phytoplankton and turbidity, but not permanently across Period 2 since there was no difference with control ponds during late summer and fall 2022 (Fig. 1, significant « Treatment × DAY_period^n^ » in models 3 and 6; Table S2).

Finally, during Period 3 corresponding to the 18 months that followed the onset of crayfish harvesting, macrophytes were not able to recover in harvested mesocosms (Fig. 1). Phytoplankton concentration was much higher in invaded mesocosms and showed marked seasonal fluctuations with peaks of concentrations over spring and summer periods (Fig. 1, significant « Treatment » and « Treatment × DAY_period^n^ » effects in model 4; Table S2). Crayfish harvesting reduced phytoplankton concentration and altered the seasonal trend with the absence of a peak in summer 2023 (Fig. 1). In 2024 however, the late summer peak was present in both harvested and invaded ponds but was weaker and ended earlier in harvested mesocosms (Fig. 1). Turbidity was higher in invaded and harvested mesocosms compared to uninvaded ones (Fig. 1, significant « Treatment » effects in model 7; Table S2), with a marked seasonal pattern characterized by summer peaks, particularly in 2024 (significant « Treatment × DAY_period^n^ »; Table S2). Alongside phytoplankton concentration, the summer turbidity peak in 2024 occurred earlier and was lower in harvested mesocosms compared to invaded ones (Fig.1).

### 2. Impacts of invasion on water temperature and ecosystem metabolism

On average, water temperature was significantly higher in both harvested and invaded mesocosms compared to uninvaded ones, but only during the warm season from approximatively May to September (i.e., significant «Treatment × DAY^n^ » effects in model 8; Table S2; Fig. 2).

**Figure 2:**
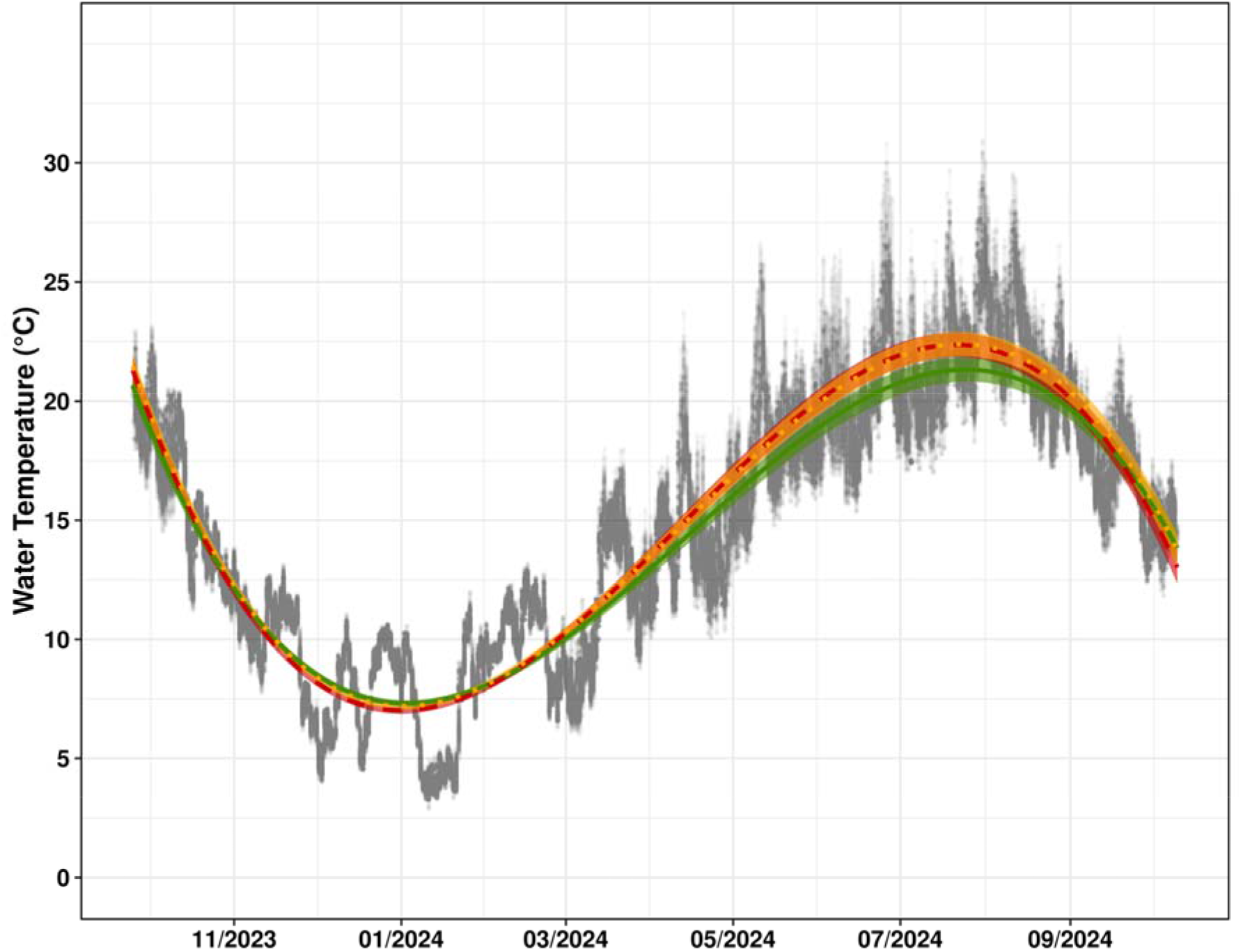
Impacts of regime shift on water temperature. Lines show LMM predictions with 95% confidence intervals for water temperature in uninvaded control (green solid line), invaded (red dashed line) and harvested mesocosms (orange dotted line) during Period 3 (Fig. 1). Grey dots show the raw data.

Experimental treatments significantly impacted the seasonal dynamics of metabolic rates (significant «Treatment × DAY^n^» effects in model 9, 10 and 11 Table S1). In uninvaded control mesocosms, net ecosystem production (NEP) decreased during spring and summer (green line in Fig. 3), reflecting a slight increase in gross primary production (GPP, green line in Fig. 3) that was overwhelmed by a strong increase in ecosystem respiration ER (green line in Fig. 3).

**Figure 3:**
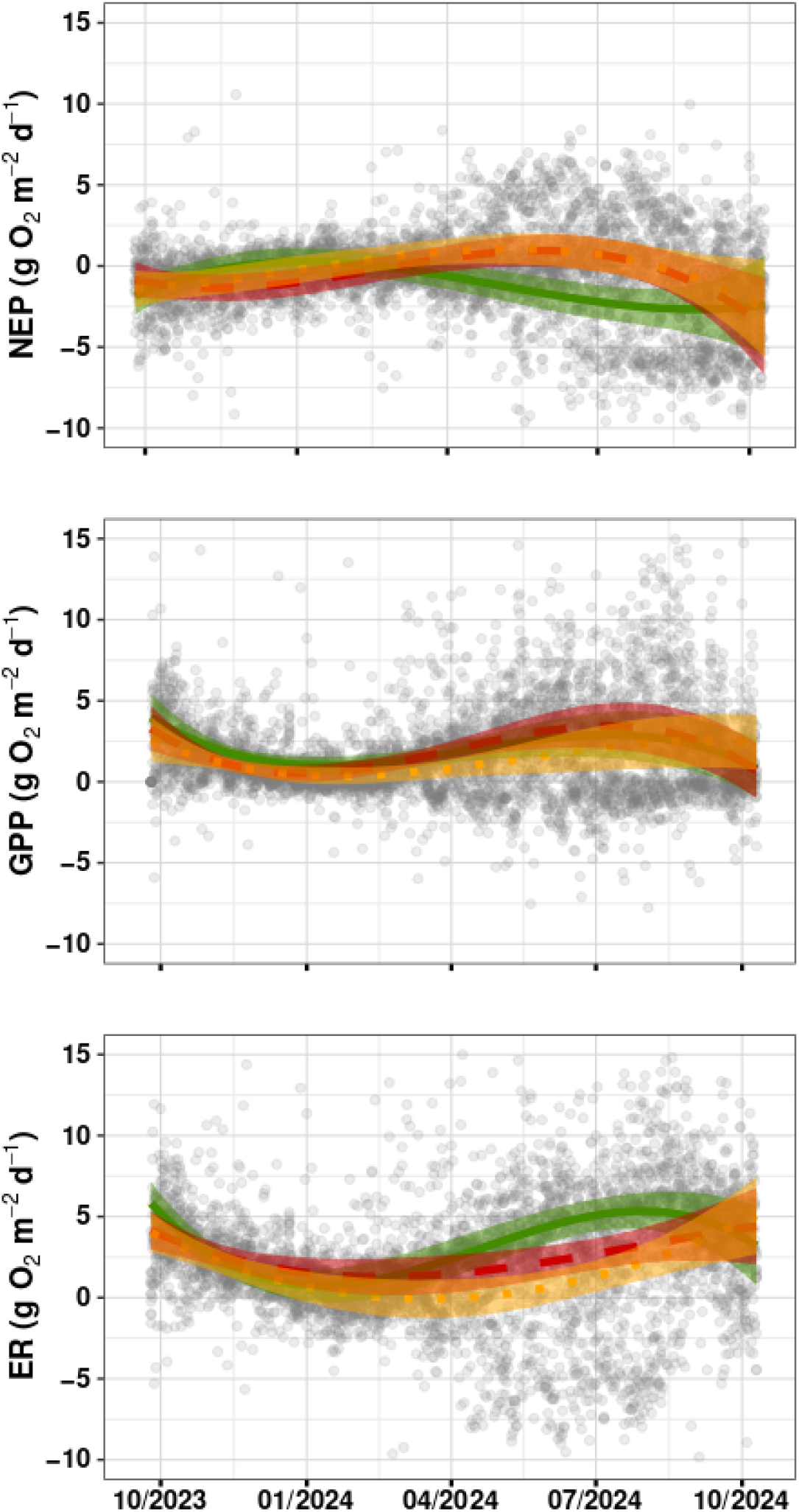
Impacts of regime shift on whole-ecosystem metabolism. Lines show LMM predictions with 95% confidence intervals for net ecosystem production (NEP), gross primary production (GPP) and ecosystem respiration (ER) in uninvaded control (green solid line), invaded (red dashed line) and harvested ponds (orange dotted line) during Period 3 (Fig. 1). Grey dots represent raw data. For clarity, we excluded extreme values from the plots but not from models (NEP: N = 15; GPP: N = 17; ER: N = 47).

In presence of crayfish, the NEP was less variable temporally, resulting in similarly increasing summer NEP in both invaded and harvested mesocosms (red and orange lines in Fig. 3, significant « Treatment × DAY^n^ » effects in model 11 Tables S1 and S2). However, this similar summer NEP increase was hiding differences in GPP and ER between invaded and harvested mesocosms. In invaded mesocosms, the summer NEP increase was driven by reduced ecosystem respiration ER only (GPP had similar seasonal patterns in uninvaded control and invaded mesocosms, green and red lines in Fig. 3). In harvested mesocosms, the summer drop in ER was stronger than in invaded mesocosms (orange and red lines in Fig. 3), but GPP also decreased significantly compared to invaded ones (orange and red lines in Fig. 3), resulting in a zero net difference on NEP.

## Discussion

Using a long-term manipulative experiment combining controlled introductions and harvesting of *P. clarkii* in pond freshwater ecosystems, we demonstrate that a single invasive species can durably drive ecosystems from a macrophyte-dominated, clear-water state to a phytoplankton-dominated, turbid state. Our findings thus provide rare experimental demonstration that an invasive species can act as a primary driver of a regime shift in ecosystems. This result challenges the prevailing view that invasive species act as “Passengers” rather than “Drivers” of ecosystem regime shifts and degradation (Dercksen *et al*. 2025). We confirm, however, previous field observations on *P. clarkii* (Rodríguez *et al*. 2003) and other reports showing that single megafaunal or invasive species can profoundly affect the structure and function of freshwater, marine or terrestrial ecosystems (Ehrenfeld 2010; Estes *et al*. 2016; He *et al*. 2024; Malhi *et al*. 2016).

Invaded ponds were pushed by crayfish toward a turbid state similarly to shallow lakes under nutrient enrichment, but the pathway to collapse differed. In classical eutrophication scenarios of regime shifts, phytoplankton grows first and precipitates macrophyte collapse, which secondarily entails sediment resuspension (Scheffer et al. 1993). We rather showed that crayfish first eradicated macrophytes and increased sediment resuspension (Fig. 1, Period 2), which entailed phytoplankton growth with some delay (Fig. 1, Period 3). This is because crayfish directly feed on macrophytes and disturb the sediments, thus removing competitors for phytoplankton and increasing nutrient availability in the water column through bioturbation and nutrient excretion (Angeler *et al*. 2001, Reynold & Aldridge 2021, Gao *et al*. 2024).

As a “press” experiment, our design does not formally demonstrate that pond mesocosms were subject to bistability (Petraitis 2013; Schröder *et al*. 2005). However, several lines of evidence suggest that they were indeed. First, control ponds maintained a very stable macrophyte-dominated, clear-water state across the five years of experiment. Second, the lack of macrophyte recovery despite crayfish harvesting, and despite sufficient time had elapsed for recovery (macrophytes initially colonized within 6 months while the harvesting period lasted 18 months), demonstrates that the transition between the two ecosystem states was nonlinear. Third, phytoplankton, a « fast » state variable due to its short generation time, did respond to crayfish harvesting by abruptly transitioning between the low and high concentrations typical of the uninvaded and invaded states, respectively (Fig. 1, Period 3). Hence, phytoplankton in harvested ponds seemingly reached a region of bistability in hysteresis, where a fast-responsive state variable can move from one to another state in response to an external forcing factor.

Modelling the drivers of this external forcing was beyond the scope of this study, but we suspect that temperature played a key role since phytoplankton concentration in harvested ponds moved from similar-to-control in winter to similar-to-invaded in spring (Fig. 1, Period 3). Such seasonal dynamics are expected in alternative stable ecosystem regimes (Scheffer & Carpenter 2003), and are consistent with the known positive effects of temperature on both phytoplankton growth and crayfish activity (Payette & McGaw 2003), and on metabolic rates in general (Brown *et al*. 2004).

Seasonal dynamics were also present in whole-ecosystem metabolism. In uninvaded control ponds, net ecosystem production (NEP) changed from null in winter to negative in summer, in line with the known heterotrophic nature of shallow lakes reported in the literature (Rabaey *et al*. 2024). In uninvaded control ponds, decreased NEP in summer resulted from increased gross primary production (GPP) being overwhelmed by an even larger increase in ecosystem respiration (ER). Such a higher sensitivity of ER than GPP to warming temperatures is predicted from the metabolic theory of ecology, and was previously observed experimentally (O’Connor *et al*. 2009, Yvon-Durocher *et al*. 2010).

Crayfish, because they increased summer temperature in invaded ponds via increased turbidity, should therefore have magnified this summer NEP decrease. However, in contrast to our expectations, we found that crayfish rather increased summer NEP to make ponds temporarily autotrophic. This result demonstrates that crayfish affected pond metabolism not only through increasing temperature, but also through other mechanisms which effects opposed and overwhelmed the effects of increasing temperature. These mechanisms are likely related to differential responses of macrophyte and phytoplankton to warming and whether the shift from macrophyte to phytoplankton dominance affected organic matter production and consumption in ponds. Although this remains a poorly-explored research area (Diamond *et al*. 2022; Hilt *et al*. 2017), we may still propose candidate mechanisms : unchanged GPP by crayfish despite higher temperatures possibly reflected a lower primary productivity of phytoplankton compared to macrophytes (Hilt *et al*. 2017), while decreased ER by crayfish despite higher temperatures possibly reflected decreased oxygenation and inhibition of microorganism respiration (Brothers *et al*. 2013). The fact that crayfish harvesting further reduced ER suggests that reduced crayfish density decreased bioturbation (as evidenced by lower turbidity in harvested mesocosms compared to invaded ones) potentially reducing microbial stimulation and associated respiration. Additionally, crayfish harvesting may have directly contributed to the decrease in ER, as crayfish metabolic activity itself represents a non-negligible component of total ecosystem respiration. Interestingly, summer autotrophy in crayfish-invaded ponds recalls the effects of nutrient enrichment, which also favours autotrophy in shallow lakes (Balmer & Downing 2011). Such a similar effect of biotic (crayfish) and abiotic (nutrient enrichment) disturbances on metabolism in aquatic systems suggests that the structural reorganization of the primary-producer community, rather than nutrient enrichment alone, drives increased autotrophy in shallow lakes worldwide.

## Conclusions

Together, our results demonstrate that a single source of biotic disturbance, an invasive species, can drive a regime shift of pond ecosystems from a macrophyte-dominated, clear-water state to a phytoplankton-dominated, turbid state. This regime shift increased water temperature in summer and changed whole-ecosystem metabolism from predominantly heterotrophic to predominantly autotrophic, indicating that invasion-induced regime shifts can interact with the seasonality of ecosystems and ramify to influence fundamental biophysical and biogeochemical processes. In the context of ongoing global change, where carbon storage is increasingly critical, our study underscores the urgent need to more broadly quantify how biological invasions alter whole-ecosystem structure, metabolism and carbon cycling.

## Acknowledgements

We are grateful to Julien Cucherousset for providing us with crayfish from Lake Lamartine. We thank Jean Marc-Paillison, Marc Collinet, Beatrice Porcon, Mingjun Feng, Eric Petit, Armand Michelot and all those who participated in the CMR field campaigns enabling the monitoring of crayfish populations. We thank Bernard Joseph, Maïra Coke and Antoine Gallard for the maintenance of mesocosms.

## Funding

This research was supported by grants from Rennes Métropole to EE (AIS18C0356 and AES1800031). A PhD grant to EG was provided by Région Bretagne (ARED 2024 CréBZH / 0985) and INRAE (ECODIV Department, 2024-9239).

## Conflicts of interests

The authors declare no conflict of interest.

## Notes

### Competing Interest Statement

The authors have declared no competing interest.

